# Comparing AI and Human Mentoring in Supporting Dental Student Well-being: A Mixed-Methods Study

**DOI:** 10.1101/2025.05.20.655059

**Authors:** Avita Rath

## Abstract

**Background:** Dental education is associated with high psychological distress among students, exacerbated by intensive clinical training and variable institutional support. Traditional mentoring programs mitigate such challenges, yet faculty shortages and resource constraints necessitate exploration of alternative models. Artificial intelligence (AI) platforms such as ChatGPT offer potential supplemental mentoring support; however, their comparative effectiveness in health professions education remains underexplored.

**Methods:** This comparative, convergent mixed-methods study was conducted at SEGi University, Malaysia, involving 31 Year 4 dental students receiving human mentoring and 36 Year 5 students utilizing ChatGPT as an AI mentor. Quantitative outcomes were assessed using the Depression, Anxiety, and Stress Scale (DASS-21), the Perceived Mentor Support Scale (PMSS), and the Mentoring Effectiveness Scale (MES). Qualitative data were collected through focus groups and analysis of ChatGPT interaction logs. Statistical analyses included paired t-tests and ANOVA, while thematic analysis was used for qualitative data integration.

**Results:** Year 4 students demonstrated significant reductions in stress, anxiety, and depression scores (p < 0.001) and greater improvements in perceived mentor support (p < 0.001) and mentoring effectiveness (p < 0.001) compared to Year 5 students. Year 5 students reported modest gains following ChatGPT mentoring, highlighting strengths in accessibility and immediacy but limitations in emotional resonance. Qualitative themes underscored human mentors’ superior capacity for emotional attunement, while ChatGPT was valued for availability and practical support.

**Conclusions:** While AI-based mentoring via ChatGPT shows promise in enhancing mentoring accessibility, human mentorship remains critical for addressing the emotional and relational needs of dental students. Careful integration of AI tools into hybrid mentorship models could optimize support systems, particularly in resource-constrained educational environments. Future research should explore ethical frameworks and training strategies to enhance empathetic AI mentorship in health professions education.

## Introduction

Dental education is one of the most demanding professional training pathways, requiring students to balance academic rigor with clinical responsibilities and patient care. This often leads to significant mental health challenges, with research indicating that up to 50% of dental students experience psychological distress during their training.^1,2^ This distress not only impacts students’ mental well-being but also has implications for their academic performance and professional readiness.^3,4^ Senior students, in particular, face heightened stress as they transition into intensive clinical practice, often without adequate institutional support.^5^

Mentoring is widely recognized as an effective strategy to mitigate stress and foster resilience in higher education. By fostering emotional well-being, resilience, and professional growth, mentoring programs significantly enhance students’ academic and personal outcomes.^6,7^ However, mentoring in dental education is often constrained by resource limitations, time constraints, and a growing student-to-faculty ratio. These challenges are especially pronounced in final-year students, who often report reduced access to mentoring when their clinical responsibilities are at their peak.^8,9^ The emergence of artificial intelligence (AI) offers a potential solution to address these mentorship gaps. Tools like ChatGPT, a conversational AI model, can simulate human-like interactions and provide real-time responses to students’ academic and emotional needs. Studies in general education and mental health contexts suggest that AI tools can improve accessibility and engagement, making them a viable alternative in resource-constrained settings.^10,11^ However, the effectiveness of AI mentoring compared to human mentoring remains underexplored, particularly in high-stress academic environments like dental education.

Recent advances in large language models (LLMs) such as ChatGPT have positioned AI as a potential supplement or substitute for academic mentoring in health professions education. Studies suggest that ChatGPT can simulate empathetic and relevant responses to educational and personal concerns when users provide sufficiently clear prompts.^12,13^However, its utility as a mentor remains underexplored in authentic educational settings, particularly in contexts where structured faculty mentoring is lacking.^14^ By comparing Year 5 dental students’ experiences with ChatGPT to Year 4 students receiving traditional mentoring, this study addresses a gap in understanding AI’s mentoring potential in resource-constrained academic environments.

This study aims to bridge this gap by evaluating ChatGPT as an AI mentor for final-year dental students at SEGi University, comparing it to traditional human mentoring provided to Year 4 students. Through a mixed-methods approach, this research will assess differences in mental health outcomes, stress levels, and satisfaction with mentoring. The findings will contribute to understanding the role of AI in educational support systems and its potential as a scalable alternative to traditional mentoring in dental education.

## Materials and Methods

### Study Design

This was a comparative, convergent mixed-methods study conducted to evaluate the effectiveness of AI mentoring (via ChatGPT) versus traditional human mentoring on the mental health, stress levels, and mentoring experiences of undergraduate dental students at SEGi University, Malaysia. The mixed-methods approach integrated quantitative and qualitative data collected simultaneously to provide a comprehensive understanding of mentoring outcomes.

### Study Setting and Population

Participants were drawn from two cohorts through convenience sampling: Year 4 and Year 5 undergraduate dental students. Thirty-one Year 4 students (5 males, 26 females) participated in monthly in-person mentoring sessions with academic staff, while 36 Year 5 students (11 males, 25 females), who did not receive scheduled academic mentoring, were introduced to ChatGPT as an AI-based mentoring tool. Inclusion criteria included full-time enrollment in the dental program and provision of informed consent. Students who were engaged in additional mentoring activities during the study period or who withdrew before completion were excluded. The sample size reflected the total number of enrolled students in each cohort during the academic term and was deemed appropriate for exploratory comparative research using mixed methods.^15^

### Ethical considerations

The study adhered to the principles of the Declaration of Helsinki.^16^ Ethical approval was obtained from SEGi University’s Ethics Committee (Approval No. SEGiEC/StR/FOD/201/2023-2024). Participants received written information about the study and its implications for policy and practice. They provided informed and written consent prior to the completion of the survey and the focus group interviews. Confidentiality and anonymity were assured, and participation was voluntary, wherein they were allowed to leave the study at any point without justification or punitive measures to their academic lives.

### Intervention

Year 4 students received structured human mentoring from faculty over a 17-week semester. Sessions were guided by the institution’s mentoring framework and addressed academic, emotional, and career-related concerns. These sessions were semi-structured, allowing for contextual flexibility.

In contrast, Year 5 students were introduced to ChatGPT and encouraged to use it as a self-directed mentoring resource. Written guidelines, including sample prompts (e.g., “How can I manage my clinical workload?”), were provided. Participants were instructed to log their interactions, which were anonymized for analysis. Students were advised that ChatGPT should be used as a supplementary tool and not as a substitute for professional counseling. Limitations, such as the tool’s lack of contextual awareness and emotional nuance, were disclosed.^13^

## Data Collection

Quantitative data were collected using three validated instruments. Mental health outcomes were assessed using the Depression, Anxiety, and Stress Scale (DASS-21), a 21-item self-report instrument measuring levels of emotional distress.^17^ Perceived mentor support was evaluated using the Perceived Mentor Support Scale (PMSS), which captures emotional and instrumental dimensions of support.^18^ Mentoring effectiveness was measured using the Mentoring Effectiveness Scale (MES), which includes relational, psychological, and communication elements.^19^

Qualitative data were gathered through separate focus group discussions for Year 4 and Year 5 participants. Each focus group included 6–8 participants and lasted approximately 60 to 90 minutes. Discussions were guided by a semi-structured interview protocol and moderated by an independent facilitator to reduce bias. All sessions were audio-recorded with participants’ consent, transcribed verbatim, and anonymized. The focus group interview guide is provided in Supporting Information (S1 Interview Guide). In addition, ChatGPT interaction logs were collected from Year 5 participants to complement focus group insights.

### Study Duration

The study was conducted over one academic semester, comprising 17 teaching weeks from 30^th^ September 2023 to 20^th^ July 2024.

## Data Analysis

Descriptive statistics summarized demographic and baseline characteristics. Within-group comparisons of pre- and post-intervention DASS-21, PMSS, and MES scores were analyzed using paired t-tests, while between-group comparisons were evaluated using one-way ANOVA. Tukey’s post-hoc test was applied where applicable, and effect sizes were reported using Cohen’s d for within-group tests and η^2^ for ANOVA.^20^

Qualitative data were analyzed using Braun and Clarke’s six-phase framework for thematic analysis.^21^ Initial coding was conducted independently by two researchers and reconciled through discussion. NVivo Qualitative Data Analysis Software version 12 (QSR International, Australia) and IBM SPSS Statistics for Windows version 26.0 were used to support analysis and ensure transparency.^22,23^

### Data Integration Strategy

This convergent mixed-methods study integrated qualitative and quantitative findings at the interpretation stage. Themes derived from focus groups and ChatGPT interaction logs were examined alongside quantitative scores on PMSS and MES to assess convergence or divergence. For instance, students’ subjective perceptions of empathy and support from ChatGPT were compared with their survey scores. While triangulation enhanced credibility, the interpretive process remained susceptible to subjectivity inherent in qualitative analysis.^24^

## Results

### Participant Characteristics

A total of 67 undergraduate dental students participated in the study, comprising 31 Year 4 students (5 males and 26 females) and 36 Year 5 students (11 males and 25 females). No significant demographic differences were observed between the two cohorts in terms of age, gender distribution, or academic performance at baseline.

### Quantitative Results

#### Mental Health Outcomes (DASS-21)

At baseline, both Year 4 and Year 5 students exhibited moderate levels of stress, anxiety, and depression, as measured by DASS-21 scores. Following the intervention, Year 4 students demonstrated a significant reduction in stress (mean difference = -4.12, *p* < 0.001, Cohen’s *d* = 0.72), anxiety (mean difference = -3.45, *p* < 0.001, *d* = 0.65), and depression scores (mean difference = -2.89, *p* = 0.002, *d* = 0.59) (Table 1). In contrast, Year 5 students also exhibited reductions in stress (mean difference = -2.11, *p* = 0.047, *d* = 0.42) and anxiety (mean difference = -1.89, *p* = 0.038, *d* = 0.39), although the magnitude of change was smaller compared to their Year 4 counterparts. No statistically significant change in depression scores was observed among Year 5 students.

**Table 1.** Changes in DASS-21 Scores Pre- and Post-Intervention.

| Group | Stress (Pre) | Stress (Post) | p-value | Anxiety (Pre) | Anxiety (Post) | p-value | Depression (Pre) | Depression (Post) | p-value |
| --- | --- | --- | --- | --- | --- | --- | --- | --- | --- |
| Year 4 (Human Mentoring) | 18.4 ± 4.2 | 14.3 ± 3.8 | <0.001 | 16.1 ± 3.9 | 12.7 ± 3.5 | <0.001 | 13.2 ± 3.5 | 10.3 ± 3.1 | 0.002 |
| Year 5 (ChatGPT Mentoring) | 17.9 ± 4.0 | 15.8 ± 3.9 | 0.047 | 15.6 ± 3.8 | 13.7 ± 3.6 | 0.038 | 13.0 ± 3.6 | 12.5 ± 3.4 | 0.158 |

*p-values calculated using paired t-tests within each group.*

#### Perceived Mentor Support (PMSS)

Year 4 students reported a significant improvement in perceived mentor support post-intervention (mean difference = +6.85, *p* < 0.001, *d* = 0.75). Year 5 students using ChatGPT also reported an increase in perceived support (mean difference = +3.21, *p* = 0.041, *d* = 0.39), albeit to a lesser extent. Between-group comparison via one-way ANOVA indicated that Year 4 students had significantly higher PMSS scores post-intervention compared to Year 5 students (*F*(1,65) = 5.72, *p* = 0.019) (Table 2).

**Table 2.** Comparison of PMSS Scores Pre- and Post-Intervention.

| Group | PMSS (Pre) | PMSS (Post) | p-value |
| --- | --- | --- | --- |
| Year 4 (Human Mentoring) | 42.1 ± 6.2 | 48.9 ± 5.8 | <0.001 |
| Year 5 (ChatGPT Mentoring) | 41.3 ± 5.9 | 44.5 ± 5.6 | 0.041 |

*p-values calculated using paired t-tests within each group. Between-group differences post-intervention analyzed using one-way ANOVA ( $p=0.019$ ).*

**Table 3:** Qualitative quotes summary.

| Theme | Exemplar Quote | Group |
| --- | --- | --- |
| Support and Connection | *“It felt like my mentor could sense when I was struggling, even when I didn’t say it outright.”* | Year 4 (Y42) |
| Support and Connection | *“When I explained my situation properly, the advice from ChatGPT felt surprisingly comforting.”* | Year 5 (Y55) |
| Responsiveness and Accessibility | *“It was like having someone ready to listen anytime, even if it wasn’t perfect.”* | Year 5 (Y53) |
| Responsiveness and Accessibility | *“It wasn’t always easy to schedule a meeting with my mentor when things got tough.”* | Year 4 (Y47) |
| Limitations and Constraints | *“ChatGPT gave textbook answers when I needed real empathy.”* | Year 5 (Y54) |
| Limitations and Constraints | *“Some mentors really cared, others just did the minimum.”* | Year 4 (Y43) |

#### Mentoring Effectiveness (MES)

Scores on the Mentoring Effectiveness Scale (MES) increased significantly for Year 4 students (mean difference = +7.34, *p* < 0.001, *d* = 0.79), particularly in relational and psychological support domains. Year 5 students demonstrated modest gains (mean difference = +2.78, *p* = 0.046, *d* = 0.38). Post-hoc analysis indicated that students receiving human mentoring perceived greater mentoring effectiveness compared to those interacting with ChatGPT (Figure 1).

**Figure 1:**
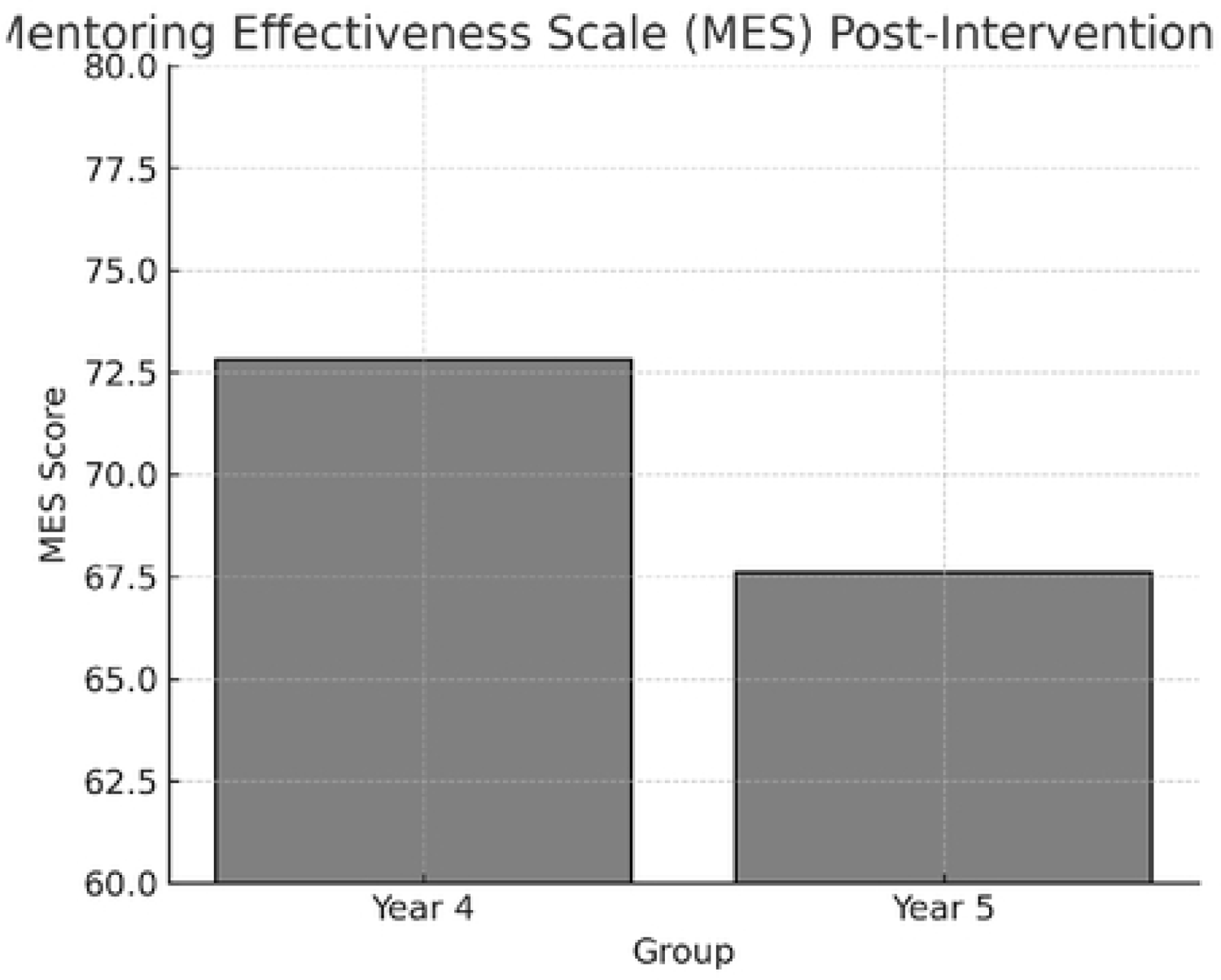
Comparison of MES Scores Pre- and Post-Intervention between years.

#### Qualitative Results

Thematic analysis of focus group discussions and ChatGPT interaction logs revealed convergent themes across both cohorts. Three primary themes emerged: **Support and Connection, Responsiveness and Accessibility**, and **Limitations and Constraints**.

#### Support and Connection

Year 4 students consistently described the value of human mentors in providing emotional reassurance and nuanced advice. One participant reflected,

> *“It felt like my mentor could sense when I was struggling, even when I didn’t say it outright.” (Y42)*

Conversely, some Year 5 students appreciated ChatGPT’s accessibility and responsiveness but acknowledged its limitations in emotional attunement. As one Year 5 student noted,

> *“ChatGPT gave good suggestions, but sometimes I needed someone who understood what being a final-year student feels like.” (Y52)*

Interestingly, a subset of Year 5 students reported that ChatGPT displayed empathetic qualities, particularly when students framed their queries thoughtfully.

> *“When I explained my situation properly, the advice felt surprisingly comforting.” (Y55)*

#### Responsiveness and Accessibility

Both groups highlighted the importance of immediate support during periods of high academic pressure. Year 5 students especially valued ChatGPT’s 24/7 availability:

> *“It was like having someone ready to listen anytime, even if it wasn’t perfect.” (Y53)*

Human mentors, while praised for depth and insight, were sometimes described as less accessible due to time constraints:

> *“It wasn’t always easy to schedule a meeting with my mentor when things got tough.” (Y47)*

#### Limitations and Constraints

Participants noted several limitations across both mentoring approaches. Some Year 5 students pointed out the superficial nature of AI-generated advice when facing complex emotional challenges:

> *“ChatGPT gave textbook answers when I needed real empathy.” (Y54)*

Meanwhile, Year 4 students discussed how mentor availability varied depending on individual faculty workload, introducing inconsistencies in mentoring quality:

> *“Some mentors really cared, others just did the minimum.” (Y43)*

#### Integration of Quantitative and Qualitative Findings

Quantitative improvements in PMSS and MES scores among Year 4 students paralleled qualitative findings highlighting stronger emotional connection and nuanced support from human mentors. For Year 5 students, the modest increases in PMSS and MES scores aligned with narratives emphasizing ChatGPT’s strengths in accessibility and informational support, but limitations in emotional resonance.

Group discussions reflected collective experiences within each cohort, particularly regarding expectations of emotional support versus practical guidance. The convergence of themes across focus groups and survey scores substantiates the reliability of the findings.

## Discussion

This study provides important insights into the role of artificial intelligence (AI)–based mentoring compared to traditional human mentoring in supporting the mental health and academic experiences of undergraduate dental students. Through a convergent mixed-methods design, it was found that while both approaches yielded improvements in stress reduction, perceived support, and mentoring effectiveness, human mentoring demonstrated significantly greater positive outcomes across all quantitative and qualitative measures. Importantly, students interacting with ChatGPT still reported notable benefits, particularly in accessibility and immediacy of responses, suggesting potential avenues for AI augmentation in mentoring structures where faculty resources are limited.

The observed reduction in DASS-21 stress and anxiety scores among students receiving human mentoring aligns with previous literature underscoring the buffering role of personalized mentoring against psychological distress in medical and dental education.^25^ These findings extend these observations by demonstrating that even a relatively modest monthly mentoring structure was sufficient to yield significant mental health benefits, supporting prior work by Gorter et al. (2008) and Neumann et al. (2011).^1,6^ In contrast, the comparatively smaller effect sizes observed among students mentored via ChatGPT suggest that while AI can offer informational support, it may lack the nuanced emotional attunement necessary for fully addressing stressors embedded within professional identity formation.^26^

Qualitative analysis further enriched these findings. Students who experienced human mentoring emphasized the relational dimension of support — noting mentors’ abilities to perceive distress even when unspoken. These findings resonate with relational mentoring theory, which posits that mentoring effectiveness is deeply rooted in mutual growth, responsiveness, and relational authenticity.^27^ In contrast, while some Year 5 students appreciated ChatGPT’s practical suggestions and empathetic phrasing when prompts were carefully crafted, others described its support as “textbook-like” and lacking deep emotional resonance — a phenomenon recently discussed in AI education research as “empathetic simulation without true sentience”.^28^

The current study also contributes to the growing discourse on the promises and perils of AI integration into educational support structures. Recent reviews by Kung et al. (2023) and Sallam (2023) highlight that large language models like ChatGPT can simulate empathy and deliver relevant advice under structured conditions, but real-world educational settings introduce variability that AI models may struggle to navigate.^12,13^ Students in my focus groups echoed this tension, acknowledging ChatGPT’s convenience while simultaneously yearning for human understanding during moments of vulnerability. These findings are consistent with recent concerns raised by Hanna and Phuong (2023) regarding the ethical design of AI systems capable of supporting emotional well-being.^29^

Notably, the results suggest that the perceived mentor support (PMSS) and mentoring effectiveness (MES) scores improved even among Year 5 students using ChatGPT, though to a lesser degree. This finding supports emerging work suggesting that AI platforms can partially bridge mentorship gaps in resource-constrained environments.^30^ However, full reliance on AI tools without supplemental human mentorship risks exacerbating feelings of isolation, particularly in clinical training settings where professional role modeling and ethical reasoning require nuanced interpersonal engagement.^10,31^

The strength of this study lies in its integrated mixed-methods design, which allowed for triangulation between quantitative outcomes and rich qualitative narratives, enhancing the credibility of findings. To my best of knowledge, this is one of the first studies to systematically compare human and AI-based mentoring using both psychometric scales and focus group analysis in a health professions education context.

Several limitations warrant consideration. First, the sample size, although reflective of the available cohorts, limits broader generalizability. Second, the self-directed nature of ChatGPT interactions may have introduced variability in engagement quality among Year 5 participants.

Third, the reliance on self-reported scales and self-logged AI interactions may introduce response bias. Future research could explore structured AI-mentoring interventions, including training students in prompt engineering to maximize empathetic responses, as suggested by Wang et al. (2023).^14^

Future directions should also include the exploration of hybrid mentoring models, wherein AI platforms are strategically integrated to complement rather than replace human mentorship. The ethical implications of expanding AI mentoring should be critically examined, including issues of data privacy, emotional authenticity, and digital equity, particularly for students from marginalized backgrounds.^32^ Finally, longitudinal studies examining the sustained effects of AI mentoring on professional identity development and well-being are warranted.

In conclusion, while ChatGPT shows promise as a supportive tool in dental education, human mentorship remains indispensable for fostering the relational, emotional, and professional development of students. Careful, ethically grounded integration of AI into mentoring ecosystems may enhance accessibility without sacrificing the essential human dimensions of academic support.

## Declarations

### Ethics Approval

Ethical approval was obtained from SEGi University’s Ethics Committee (SEGiEC/StR/FOD/201/2024-2025). Written informed consent was obtained from all participants.

### Data Availability

The datasets generated and/or analyzed during the current study are not publicly available due to institutional policies regarding participant confidentiality but are available from the corresponding author on reasonable request.

